# HViLM: A Foundation Model for Viral Genomics Enables Multi-Task Prediction of Pathogenicity, Transmissibility, and Host Tropism

**DOI:** 10.64898/2026.03.18.712700

**Authors:** Pratik Dutta, Jack Vaska, Pallavi Surana, Rekha Sathian, Max Chao, Zhihan Zhou, Han Liu, Ramana V. Davuluri

**Affiliations:** Department of Biomedical Informatics, Stony Brook University, Stony Brook, NY 11794; Department of Computer Science, Northwestern University, Evanston, IL 60208, USA

## Abstract

**Motivation:** The emergence of novel viral pathogens poses critical threats to global health, yet current computational approaches for viral risk assessment are predominantly virus-specific and require extensive retraining for each new threat. Computational methods for rapid characterization of emerging viruses across multiple epidemiologically relevant dimensions—pathogenicity, host tropism, and transmissibility—are urgently needed to inform public health responses and guide experimental prioritization.

**Results:** We present **HViLM (Human Virome Language Model)**, the first foundation model for panviral genomic analysis through continued pre-training of DNABERT-2 on 5 million non-redundant viral sequences (MMseqs2-clustered from 25 million chunks at 80% identity) spanning 9,000 species across 45+ viral families from the VIRION database. We introduce the **Human Virome Understanding Evaluation (HVUE) benchmark** comprising seven curated datasets across three prediction tasks: pathogenicity classification, host tropism prediction, and transmissibility assessment. Through parameter-efficient fine-tuning with LoRA, HViLM achieves state-of-the-art performance with average accuracies of 95.32% for pathogenicity, 96.25% for host tropism, and 97.36% for transmissibility assessment. The model demonstrates robust cross-family generalization, substantially outperforming sequence-similarity baselines and general genomic foundation models. Attention-based interpretability analysis reveals that HViLM captures biologically meaningful pathogenicity determinants through molecular mimicry of host regulatory elements, including convergent evolution of eight independent sequences targeting Interferon Regulatory Factor 1 (Irf1) for immune evasion.

**Availability:** The HVUE benchmark datasets, training scripts, and complete implementation are publicly available at https://github.com/duttaprat/HViLM. Pre-trained HViLM-base model weights and fine-tuned task-specific variants are available on Hugging Face at https://huggingface.co/duttaprat/HViLM-base.

**Supplementary information:** Supplementary data are available online.

## 1 Introduction

The emergence of novel viral pathogens poses critical threats to global health, yet current computational approaches for viral risk assessment are predominantly virus-specific and require extensive retraining for each new threat. Traditional methods rely on sequence alignment (BLAST(Koltes, et al., 2009), HMMER(Potter, et al., 2018)) or k-mer-based machine learning classifiers that struggle with computational efficiency, sensitivity to novel pathogens, and generalization across viral families. The COVID-19 pandemic underscored the urgent need for rapid computational characterization of emerging viruses across multiple epidemiologically relevant dimensions—pathogenicity (disease-causing potential), host tropism (species infectivity), and transmissibility (epidemic potential)—to inform public health responses and guide experimental prioritization.

Recent advances in genomic foundation models such as DNABERT(Ji, et al., 2021), DNABERT-2(Zhou, et al., 2023), and Nucleotide Transformer(Dalla-Torre, et al., 2025) leverage self-supervised learning on millions of genomic sequences to learn contextual representations that generalize across species and biological contexts. However, existing models are pre-trained primarily on prokaryotic genomes with limited viral representation and have been applied predominantly to single-task scenarios, lacking comprehensive benchmarks for multi-task viral phenotype prediction essential for pandemic preparedness.

Recent work has explored foundation models for pathogen detection. PathoLM(Dip, et al., 2024) pioneered genomic language models for bacterial pathogenicity prediction, demonstrating superior performance through transfer learning from Nucleotide Transformer(Dalla-Torre, et al., 2025), but focused exclusively on bacterial pathogens and single-task classification. Recent viral-specific models have emerged but face limitations: ViraLM(Peng, et al., 2024) focused primarily on taxonomic classification rather than epidemiologically relevant phenotypes, while VirRep(Dong, et al., 2024) achieved strong viral family classification through contrastive learning but lacks interpretability mechanisms and multi-task evaluation. Unlike prior viral genomics models that function as black-box classifiers or focus on single tasks or limited viral families, HViLM jointly introduces (i) ***large-scale viral-specialized continued pretraining***, (ii) ***a unified multi-task evaluation benchmark (HVUE)***, and (iii) ***a systematic interpretability framework linking sequence representations to host regulatory mimicry***.

We address this gap by presenting **HViLM (Human Virome Language Model)**, the first foundation model for comprehensive viral risk assessment through multi-task prediction of pathogenicity, host tropism, and transmissibility:

1. **Viral-Specialized Foundation Model**: We develop HViLM by continued pre-training of DNABERT-2 on 5 million curated viral genome sequences spanning 45+ viral families from the VIRION database(Carlson, et al., 2022), specifically optimizing the model to capture viral genomic patterns relevant for human disease risk assessment.
2. **HVUE Benchmark:** We introduce the Human Virome Understanding Evaluation (HVUE) benchmark, comprising seven carefully curated datasets totaling 220,000 viral sequences across three critical prediction tasks: pathogenicity classification (distinguishing disease-causing from benign strains), host tropism prediction (identifying human-infecting viruses), and transmissibility assessment (evaluating epidemic potential via R_0_-based classification(Obadia, et al., 2012)). HVUE provides the first systematic multi-task evaluation framework for viral genomics foundation models.
3. **Multi-Task Predictive Framework:** Through parameter-efficient Low-Rank Adaptation (LoRA) (Hu, et al., 2022) fine-tuning, HViLM achieves state-of-the-art performance across all tasks (95.32% pathogenicity, 96.25% host tropism, 97.36% transmissibility), substantially outperforming sequence alignment baselines and general genomic foundation models.
4. **Mechanistic Interpretability:** Attention-based analysis reveals that HViLM learns biologically meaningful pathogenicity determinants through molecular mimicry of host regulatory elements. We identify 42 conserved motifs matching 10 vertebrate transcription factors, with convergent evolution of 8 independent sequences targeting Interferon Regulatory Factor 1 (Irf1) for immune evasion(Myoung, et al., 2019) and multiple motifs targeting Foxq1 for epithelial tropism(Chetta, et al., 2025; Feuerborn, et al., 2011), demonstrating coordinated viral strategies to hijack host regulatory machinery.

## 2 Datasets

### 2.1 Pre-training Dataset

HViLM was pre-trained on viral genomes from the VIRION database(Carlson, et al., 2022), a comprehensive collection of host-virus interactions maintained by Verena (Viral Emergence Research Initiative), encompassing 476,242 documented virus-host associations across 9,000 viral species and 3,767 vertebrate host species. From 10,817,265 unique NCBI accession numbers, we retrieved complete viral genome sequences and segmented them into non-overlapping 1000 base pair chunks. To remove redundancy while preserving viral diversity, we applied MMseqs2(Mirdita, et al., 2019) sequence clustering at 80% identity threshold, reducing the dataset to 5 million unique sequences representing distinct viral genomic regions across 45+ viral families spanning all Baltimore classification groups.

#### 2.1.1 Large-Scale Sequence Retrieval and Quality Control Pipeline

We constructed a large-scale viral genome pre-training corpus using the VIRION (Viral Emergence Research Initiative Online) database, a comprehensive, manually curated resource maintained by Verena that integrates virus–host associations from GenBank, peer-reviewed literature, outbreak investigations, and experimental infection studies. VIRION provides broad vertebrate host coverage, encompassing approximately 25% of known mammalian diversity, 10% of avian diversity, and 6% of total vertebrate diversity, including key viral reservoir taxa such as bats, rodents, primates, and domestic species. All viral and host records undergo full taxonomic reconciliation against the NCBI Taxonomy database, resolving synonymy and historical nomenclature inconsistencies. The curated resource includes ~9,000 validated viral species across major RNA and DNA virus families and 3,767 vertebrate host species, forming one of the most comprehensive reconciled virus–host datasets available for large-scale computational modeling.

From 476,242 VIRION-documented virus–host interaction records, we retrieved complete viral nucleotide sequences using a scalable, parallelized NCBI Entrez Direct pipeline, yielding ~187 GB of raw FASTA data. Retrieved sequences were subjected to multi-stage quality control, including removal of short fragments (<500 bp) and exact duplicates arising from redundant submissions or technical replicates. Each quality-controlled viral genome was segmented into non-overlapping 1,000 bp chunks, discarding terminal fragments shorter than 500 bp, generating ~25 million training sequences. To reduce redundancy while preserving viral diversity, we applied MMseqs2 clustering at 80% sequence identity and 80% coverage, collapsing closely related strains and reducing the dataset five-fold to ~5 million representative sequences. This threshold reflects standard viral genomics practice and preserves species- and subspecies-level diversity. The resulting corpus provides a high-quality, diversity-preserving foundation for pre-training genome-scale language models capable of capturing viral sequence structure across broad evolutionary and ecological contexts.

### 2.2 Human Virome Understanding Evaluation (HVUE) Benchmark

We introduce the Human Virome Understanding Evaluation (HVUE) benchmark, consisting of seven curated datasets from NCBI Virus, BV-BRC(Shukla, et al., 2025), and Virus-Host DB(Mihara, et al., 2016), spanning three prediction tasks critical for pandemic preparedness: pathogenicity classification, host tropism prediction, and transmissibility assessment. HVUE comprises seven curated datasets from NCBI Virus, BV-BRC, and Virus-Host DB, totaling 220,000 viral sequences spanning 30+ viral families and generating 900,000+ standardized sequence fragments for model training (Table 1). To rigorously evaluate cross-family generalization—critical for pandemic preparedness where novel threats may emerge from previously uncharacterized families—datasets provide both single-family deep coverage (BVBRC-CoV with 18K sequences, BVBRC-Calici with 31K sequences, Orthomyxoviridae with 152K sequences) and broad phylogenetic diversity (CINI spanning 4 families, VHDB spanning 30 families).

**Table 1.**
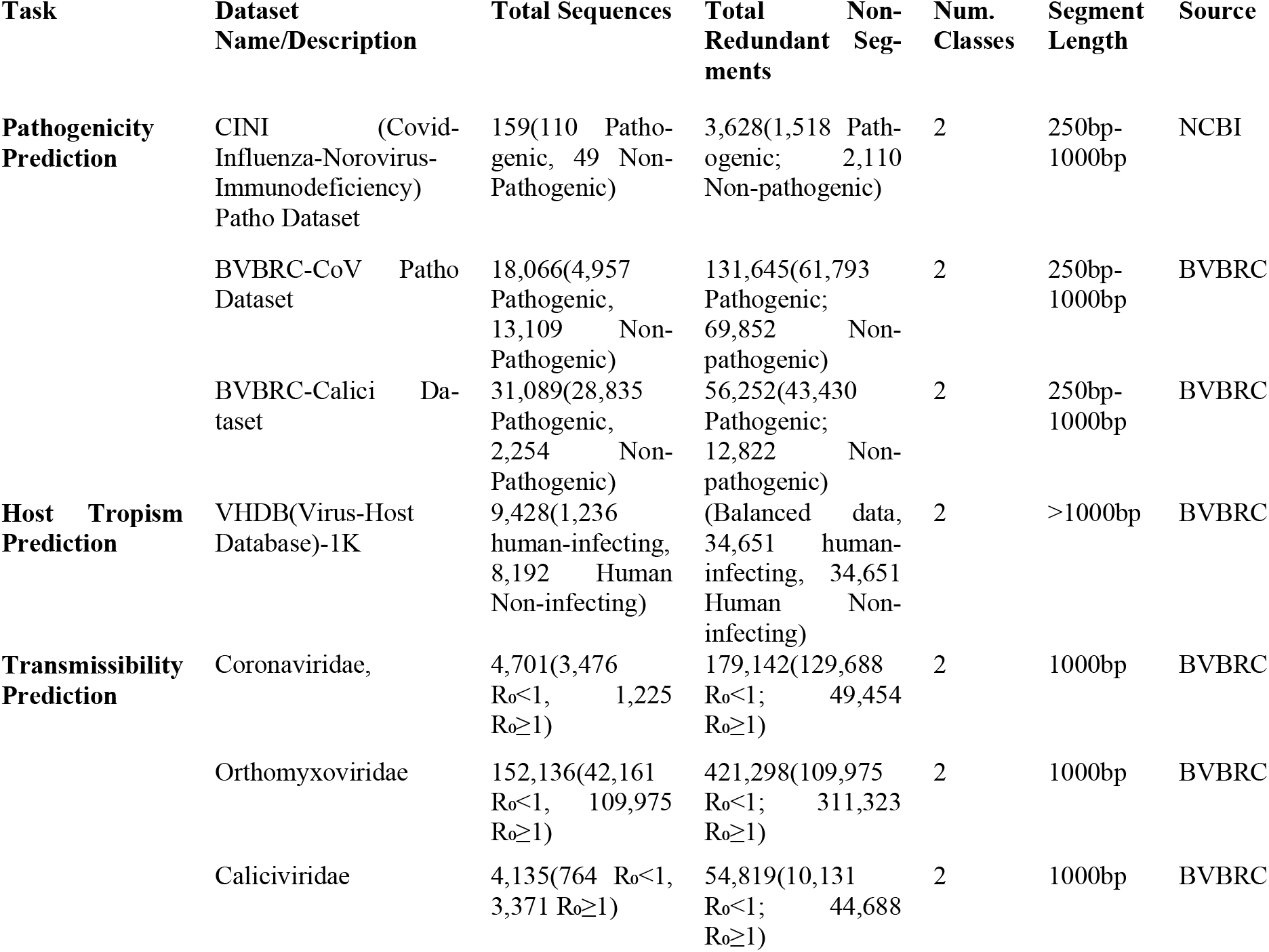
HVUE Benchmark Datasets Summary. Overview of all Human Virome Understanding Evaluation (HVUE) benchmark datasets, including prediction task, dataset description, viral families covered, total genome sequences, non-redundant sequence segments after fragmentation (250–1000 bp), number of classes, segment length, and data sources. For transmissibility prediction, genome-level R_0_ annotations were derived from epidemiological literature and sequences were segmented into 1000 bp fragments for model training. Balanced (1:1) splits were used for host tropism evaluation.

#### Pathogenicity Prediction (3 datasets)

CINI (159 sequences, 4 viral families with manual literature-based curation), BVBRC-CoV (18,066 coronaviruses), and BVBRC-Calici (31,089 caliciviruses) yielding 191,525 training segments. Labels distinguish viruses causing human disease from non-pathogenic strains based on clinical evidence and isolation sources, following pathogenicity classification criteria adapted from PathoLM and refined through systematic PubMed and ViralZone literature searches (Supplementary Table S1).

#### Host Tropism Prediction (1 dataset)

VHDB dataset (9,428 sequences, 30 viral families) with experimentally validated host range annotations, formulated as binary classification of human-tropic (n=1,236, 13.1%) vs. non-human-tropic (n=8,192, 86.9%) viruses. While the original dataset exhibits 1:6.6 class imbalance reflecting biological reality—the vast majority of viral diversity exists in non-human reservoirs—we applied balanced (1:1) sampling for model training to prevent overwhelming bias toward the majority class. This design requires models to identify subtle molecular adaptations enabling human infection (receptor binding specificity, replication compatibility, immune evasion) rather than exploiting class distribution shortcuts. Sequences were segmented into fragments >1000 bp to capture complete open reading frames across phylogenetically diverse viral families.

#### Transmissibility Prediction (3 datasets)

Three family-specific R_0_-based binary classification datasets (R_0_ < 1 vs. R_0_ ≥ 1): Coronaviridae (4,701 sequences, 179,142 segments), Orthomyxoviridae (152,136 sequences, 421,298 segments), and Caliciviridae (4,135 sequences, 54,819 segments) based on documented basic reproduction numbers from epidemiological literature. The R_0_ = 1 epidemic threshold separates viruses capable of sustaining human transmission chains (e.g., SARS-CoV-2 R_0_=2.0-3.0, seasonal influenza R_0_=1.3) from those causing only sporadic spillover events without sustained transmission (e.g., MERS-CoV R_0_=0.3-0.8, H5N1 R_0_<0.5). Sequences were segmented into fixed 1000 bp windows for consistent model training. Complete R_0_ assignments with primary literature citations are provided in Supplementary Table S2.

Pathogenicity datasets were segmented into variable-length 250-1000 bp non-overlapping fragments to handle diverse genome sizes across viral families while preserving sufficient genomic context for pathogenicity-associated features. This window size captures complete functional domains including receptor binding domains (typically 400-600 bp), packaging signals, and regulatory elements, while enabling consistent model training across diverse genome architectures ranging from small RNA viruses (~7-12 kb) to large DNA viruses (>100 kb). Non-overlapping segmentation ensures statistical independence between training instances and prevents information leakage across train-validation-test splits.

### 2.3 Data Leakage Prevention and Dataset Splitting

To ensure fair evaluation and prevent information leakage, we performed systematic overlap analysis between the VIRION pre-training corpus (10.8M sequences) and HVUE benchmark datasets (58K+ sequences) through accession ID matching and MMseqs2 similarity analysis at >95% identity threshold. We identified and excluded 186 overlapping sequences from the pre-training corpus before model training began. Following decontamination, all HVUE datasets were split into training (70%), validation (15%), and test (15%) sets using stratified sampling to maintain class distribution across pathogenicity labels and viral family representation.

## 3 Methods

### 3.1 Model Architecture and Pretraining

HViLM builds upon DNABERT-2, a genomic foundation model based on the MosaicBERT(Portes, et al., 2023) architecture pretrained on 135 million prokaryotic and viral genomes. DNABERT-2 employs BPE tokenization and processes sequences up to 1000 base pairs through a 12-layer transformer encoder with 117 million parameters (768 hidden dimensions, 12 attention heads). We initialize HViLM with DNABERT-2’s pretrained weights, leveraging its learned representations of fundamental genomic patterns, including codon usage bias, GC content variation, regulatory motifs, and evolutionary conservation—to enable effective transfer learning for viral-specific prediction tasks.

To enhance the model’s capability for viral-specific genomic pattern recognition, we performed continued pre-training (domain-adaptive pre-training)(Gururangan, et al., 2020) on 5 million curated viral genome sequence chunks derived from the VIRION database (Section 2.1). As illustrated in Figure 1, the pre-training pipeline begins with 10.8 million complete viral genomes from VIRION spanning 9,000 viral species and 3,767 vertebrate hosts. Following quality control filtering and 1000bp chunking, we applied MMseqs2 sequence clustering at 80% identity threshold to remove phylogenetic redundancy while preserving species- and subspecies-level diversity, yielding 5 million representative chunks covering 45+ viral families.

**Figure 1.**
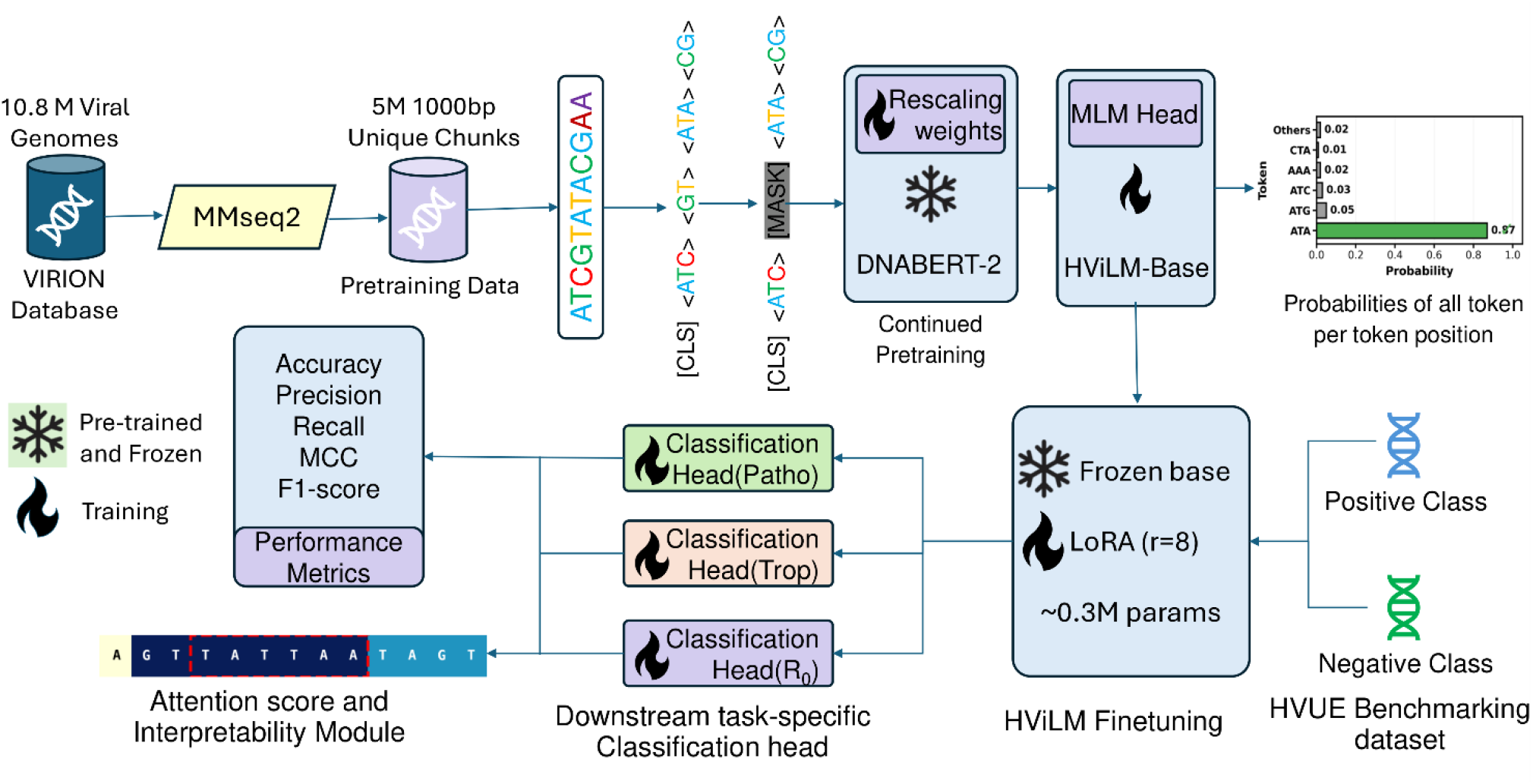
Overview of HViLM Pipeline for Multi-Task Viral Risk Assessment. HViLM employs a three-stage approach: (1) Continued pre-training of DNABERT-2 on 5M viral sequences from VIRION database with masked language modeling, (2) Parameter-efficient fine-tuning using LoRA on HVUE benchmark datasets for three prediction tasks (pathogenicity, host tropism, transmissibility), and (3) Interpretability analysis through attention-guided motif discovery revealing transcription factor mimicry mechanisms.

This continued pre-training employed masked language modeling (MLM) objective, where 15% of input tokens were randomly masked and the model learned to predict the original tokens based on bidirectional context (Figure: - 1). where 15% of input BPE tokens were randomly selected for masking using a three-way strategy: (i) 80% of selected tokens replaced with the special [MASK] token, (ii) 10% replaced with random tokens sampled from the vocabulary to introduce noise robustness, and (iii) 10% left unchanged to reduce pre-training-to-fine-tuning mismatch. The model learns to predict original masked tokens based on bidirectional context from surrounding sequences, enabling it to capture both local sequence patterns (e.g., start/stop codons, ribosomal binding sites, splice junctions) and long-range dependencies (e.g., RNA secondary structure constraints, inter-domain interactions, regulatory element spacing) specific to viral genome organization.

Training was conducted for 10 epochs using AdamW optimizer(Loshchilov and Hutter, 2016) with learning rate 5e-5, linear warmup over 10% of training steps, and cosine decay schedule. We used batch size 32 with gradient accumulation over 4 steps (effective batch size 128) and mixed precision training (fp16) on 4 NVIDIA A100 GPUs. Training converged after approximately 72 hours (288 GPU-hours total), achieving 94.2% MLM accuracy on a held-out validation set of 500,000 viral sequence chunks (10% of training data) not present in the pre-training corpus. Convergence was determined by monitoring validation loss, which plateaued within 0.5% for three consecutive epochs, indicating the model had learned stable representations without overfitting. This viral-specialized base model serves as the foundation for all downstream task-specific fine-tuning.

### 3.2 Task-Specific Fine-Tuning

From the viral-specialized HViLM base model, we developed three task-specific variants through parameterefficient fine-tuning. (1) **HViLM-Patho**: Fine-tuned for pathogenicity classification to distinguish diseasecausing from non-pathogenic viral strains across three datasets (CINI, BVBRC-CoV, BVBRC-Calici) totaling over 50,000 segmented viral sequences. (2) **HViLM-Tropism**: Fine-tuned for host tropism prediction to classify human-tropic versus non-human-tropic viruses on 9,428 complete viral genomes spanning 30 viral families from the VHDB dataset. (3) **HViLM-R0**: Fine-tuned for transmissibility assessment to predict epidemic potential through R_0_-based(Delamater, et al., 2019) binary classification (R_0_<1 vs R_0_≥1) across three viral familyspecific datasets (Coronaviridae, Orthomyxoviridae, Caliciviridae).

To efficiently adapt the 117M-parameter base model while avoiding catastrophic forgetting of pre-trained knowledge and reducing computational requirements, we employed Low-Rank Adaptation (LoRA)(Hu, et al., 2022). LoRA freezes the pre-trained model weights and injects trainable low-rank decomposition matrices into the attention layers, enabling task-specific adaptation with minimal additional parameters. We applied LoRA to the query and value projection matrices in all 12 attention layers with rank r=8 and scaling factor α=16. This configuration introduces only ~0.3 million trainable parameters (~0.26% of total model parameters) per task while maintaining model expressiveness. For each task, we added a classification head comprising dropout (p=0.1) and a linear layer mapping [CLS] representations to binary outputs.

**Table 2.** Performance comparison of HViLM variants and baseline genomic foundation models on HVUE benchmark tasks. All models were fine-tuned using identical LoRA-based procedures on the same data splits. Metrics computed on held-out test sets. Bold indicates best performance per dataset. HViLM variants consistently outperform baselines across pathogenicity classification, host tropism prediction, and transmissibility assessment tasks.

Hyperparameters were systematically optimized using Optuna(Akiba, et al., 2019) with Tree-structured Parzen Estimator algorithm on validation sets. Models were trained on NVIDIA A100 GPUs with mixed precision training, using stratified sampling and early stopping with patience of 3 epochs.

### 3.3 Comparative Genomic Foundational Models

We compared HViLM against state-of-the-art genomic foundation models to contextualize performance gains from viral-specific continued pre-training. Baseline models include: (1) Nucleotide Transformer 500M-1000g (NT-500M-1000g)(Dalla-Torre, et al., 2025), pre-trained on DNA sequences from over 3,200+ diverse human genomes; (2) GENA-LM(Fishman, et al., 2025), a transformer-based foundational DNA language model pre-trained on multi-species genomes with 36kb context; and (3) DNABERT-MB(Jack Vaska, 2025), pretrained on human microbiome-derived genomic sequences. All comparative models were fine-tuned using the same training procedures as HViLM, including LoRA-based adaptation and task-specific classification heads. All models were fine-tuned using identical LoRA procedures and evaluated on the same data splits, isolating the contribution of viral-specific pre-training.

### 3.4 Evaluation Metrics

We assessed all the HViLM Finetuned models and comparative genomic foundation models Model performance using metrics suited for binary classification under class imbalance, including accuracy, precision, recall, F1-score, and Matthews Correlation Coefficient (MCC). All metrics were computed on held-out test sets using predictions from the best-performing model checkpoint (highest validation F1-score).

### 3.5 Computational Infrastructure and Efficiency

Continued pre-training on 5 million viral chunks was performed on 4 NVIDIA A100 80GB GPUs using mixed precision training (fp16), completing in approximately 72 hours (288 GPU-hours total). The training employed AdamW optimizer with learning rate 5e-5, batch size 32 with gradient accumulation over 4 steps (effective batch size 128), achieving 94.2% MLM accuracy on held-out validation sequences.

Task-specific fine-tuning via LoRA (rank r=8, scaling α=16) introduced only ~0.3 million trainable parameters (~0.26% of DNABERT-2’s 117M parameters) per task, enabling adaptation in <6 hours per task on a single NVIDIA A100 GPU. This parameter efficiency makes HViLM practical for rapid deployment during emerging infectious disease outbreaks where computational resources may be limited but fast characterization is critical.

Compared to training task-specific models from scratch (estimated 200-300 GPU-hours per task based on similar-scale architectures), our transfer learning approach achieves 30-50× computational savings while maintaining superior performance, demonstrating the practical value of foundation models for pandemic preparedness applications.

## 4 Results

### 4.1 Pathogenicity Classification Performance

HViLM-Patho demonstrated superior performance across all three pathogenicity datasets, achieving average accuracies of 95.32% (NT-500M-1000g), 89.97% (GENA-LM), and 92.73% (DNABERT-MB; Table 1). On the manually curated CINI dataset, which includes diverse viral families with literature-validated pathogenicity labels, HViLM-Patho achieved the strongest performance with balanced metrics across both pathogenic and non-pathogenic classes (MCC: 74.48). For the larger BVBRC-CoV dataset, HViLM-Patho achieved 98.26% accuracy with remarkably balanced precision (98.23%) and recall (98.28%), demonstrating robust discrimination between human-pathogenic coronaviruses (SARS-CoV-2, SARS-CoV, MERS-CoV, seasonal HCoVs) and animal-restricted strains. The highly imbalanced BVBRC-Calici dataset showed exceptional performance across all models due to strong class separability, though HViLM-Patho maintained the highest metrics across all measures. Notably, viral-specific continued pre-training provided consistent advantages over general genomic foundation models, with improvements most pronounced on the challenging CINI dataset where HViLM-Patho outperformed DNABERT-MB by 5.7 percentage points.

### 4.2 Host Tropism Prediction Performance

HViLM-Tropism achieved 96.25% accuracy on the VHDB dataset for distinguishing human-tropic from nonhuman-tropic viruses across 30 viral families (Table 1). Our model’s performance was comparable to DNABERT-MB (96.39%) but substantially exceeded NT-500M-1000g and GENA-LM, which showed approximately 10 percentage point drops. The similar performance between HViLM-Tropism and DNABERT-MB is notable given that microbiome-specific pre-training captures bacteria-virus host interactions relevant to tropism prediction.

The model demonstrated strong generalization across diverse taxonomic groups including RNA viruses (Caliciviridae, Picornaviridae, Flaviviridae), DNA viruses (Papillomaviridae, Herpesviridae), and retroviruses, indicating that learned representations capture universal sequence features associated with human infectivity rather than family-specific patterns. The substantial gap between domain-specific models (HViLM, DNABERT-MB) and general genomic models demonstrates that pre-training on biologically related sequences, whether viral or microbiome-focused, enhances host tropism prediction.

### 4.3 Transmissibility Assessment Performance

HViLM-R0 achieved consistently high performance across all three viral families for R_0_-based transmissibility classification, with an average accuracy of 97.36% (Table 1). For Coronaviridae, HViLM-R0 correctly classified human-transmissible coronaviruses capable of epidemic spread (SARS-CoV-2 variants, SARS-CoV, seasonal HCoVs) while identifying subcritical viruses like MERS-CoV and bat coronaviruses. Caliciviridae showed near-perfect classification, distinguishing highly transmissible human noroviruses from animal-restricted caliciviruses.

Compared to baselines, HViLM-R0 showed modest but consistent improvements over DNABERT-MB across all families, with larger advantages over NT-500M-1000g and GENA-LM. Notably, GENA-LM exhibited highly variable performance across families (ranging from 81.66% to 93.73%), while HViLM-R0 maintained stable high performance (95.62-99.01%), suggesting that viral-specific pre-training enables learning of universal transmissibility-associated genomic patterns that generalize across viral families with distinct transmission biology.

### 4.5 Interpretability Analysis Through Attention Mechanisms

To validate that HViLM learns biologically meaningful features rather than spurious correlations, we performed attention analysis on BVBRC-CoV pathogenic and non-pathogenic coronaviruses. Pathogenic sequences exhibited significantly lower mean attention scores but higher variance (0.032 vs 0.045, p=3.45×10^−14^), indicating focused attention on specific genomic regions rather than diffuse sequence scanning (Supplementary Figure S1A-B). We extracted high-attention genomic regions (attention scores exceeding the 95th percentile across all sequences) and applied MEME-ChIP motif discovery to identify conserved sequence patterns. This analysis revealed 42 distinct motifs ranging from 14 to 20 base pairs that were consistently associated with elevated attention scores in pathogenic viral sequences (Supplementary Table S3). To determine whether these attention-focused motifs represent functional regulatory elements, we performed TOMTOM matching against the JASPAR(Castro-Mondragon, et al., 2022) vertebrate transcription factor binding site database. Strikingly, the discovered viral motifs exhibited significant sequence similarity to host transcription factor binding sites, suggesting molecular mimicry as a core mechanism underlying coronavirus pathogenicity.

The most striking pattern was convergent evolution toward host transcription factor mimicry. Eight independent viral motifs—TTTTATTA, TTATTAAA, TATTAA, TATTA, ATTTATT, TTATTAA, ATATTTT, and AATTTAT—all matched Interferon Regulatory Factor 1 (Irf1)(Reis, et al., 1992; Taniguchi, et al., 2001) binding sites (JASPAR MA0050.4), with local enrichment ratios of 1.02-1.13× and p-values from 4.5×10^−4^ to 4.0×10^−3^. This redundancy indicates strong positive selection for Irf1 mimicry across coronavirus genomes(Lei, et al., 2020). Similarly, three motifs (AAAAAATT, TATAAAAA, TATAAA) matched Foxq1 binding sites(Feuerborn, et al., 2011), with AAAAAATT showing the strongest enrichment (1.20× local, p=3.9×10^−10^; 1.19× global, p=9.5×10^−7^). Additional matches included ZNF354A (6 motifs), BARHL2 (5 motifs), and BARX2, LIN54, FOXD3, and Pgr (4 motifs each), totaling 10 unique transcription factor targets across 42 viral motifs (complete mappings in Supplementary Table S3).

Representative examples illustrate precise molecular mimicry (Figure 2). AAAAAATT exhibited the strongest statistical enrichment (1.20× local, p=3.9×10^−10^) and matched Foxq1 binding sites (10 bp overlap), a forkhead box transcription factor regulating epithelial differentiation in respiratory tissues—the primary coronavirus entry site. TATTAA demonstrated high global enrichment (1.27×, p=9.9×10^−8^) and matched Irf1 (11 bp overlap), the master regulator of interferon-mediated immunity. TAATAA showed significant enrichment (1.10× local, p=2.0×10^−4^; 1.15× global, p=3.9×10^−3^) with exceptional ZNF354A match precision (19 bp overlap, q=0.056). TTTTATTA exhibited robust enrichment (1.10× local, p=4.0×10^−3^; 1.14× global, p=1.6×10^−3^) and also matched Irf1, visually demonstrating convergent evolution where sequence-distinct motifs independently target the same host immune regulator. The redundancy of immune evasion mechanisms is visually apparent in Figure 2B and 2D, where two sequence-distinct motifs (TATTAA and TTTTATTA, sharing only 50% sequence identity) independently converged on Irf1 binding site mimicry, demonstrating strong positive selection for interferon response suppression.

**Figure 2:**
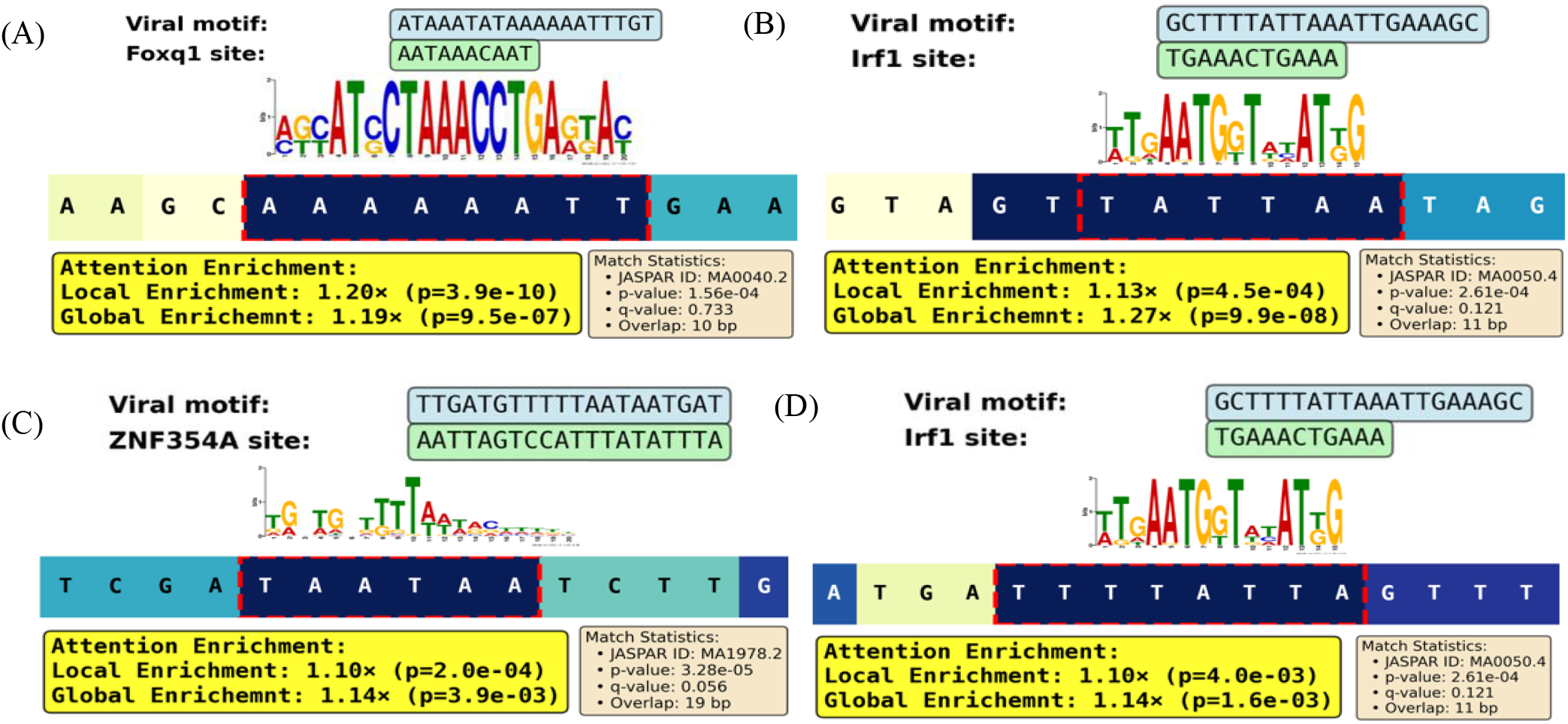
HViLM Identifies Host Transcription Factor Binding Site Mimicry Through Attention-Guided Motif Discovery in Pathogenic Coronaviruses. (A) AAAAAATT matches Foxq1 (1.20× local enrichment, p=3.9×10^−10^), regulating respiratory epithelial tropism. **(B)** TATTAA matches Irf1 (1.27× global, p=9.9×10^−8^), controlling interferon responses. **(C)** TAATAA matches ZNF354A (19 bp overlap, q=0.056), regulating immune pathways. **(D)** TTTTATTA motif shows robust enrichment and also matches Irf1 binding site, demonstrating convergent evolution of independent sequences for immune evasion. Dark blue shading indicates high-attention regions; sequence logos show MEME-derived viral motifs aligned with JASPAR vertebrate transcription factor consensus sequences.

These findings establish that HViLM captures genuine pathogenicity determinants through host regulatory mimicry. The mechanistic coherence—Foxq1 mimicry for epithelial entry, Irf1 mimicry for immune evasion (demonstrated by 8 convergent motifs), and ZNF354A mimicry for chromatin regulation—reveals coordinated multi-target genomic strategies. The convergent evolution of 8 distinct Irf1-matching motifs indicates immune suppression represents the strongest selective pressure, with redundancy providing mutational robustness. Unlike alignment or k-mer methods that miss dispersed non-homologous motifs, HViLM’s transformer architecture automatically learns these functional elements through attention mechanisms, enabling both accurate prediction and mechanistic discovery of therapeutic targets.

### 4.5 Systematic Interpretability Framework for Pathogenicity Determinants

Beyond the discovered motifs, we developed a comprehensive computational framework to systematically identify and validate biologically meaningful pathogenicity determinants through attention-guided motif discovery. Our four-stage pipeline:

1. **Token-Level Attention Analysis:** For each viral sequence, we extracted attention weights from HViLM’s final transformer layer (layer 12) to identify genomic regions with elevated attention scores (>95th percentile) during pathogenicity prediction.
2. **Motif Discovery and Consensus Generation:** High-attention tokens were clustered by sequence similarity using MEME-ChIP, yielding 42 distinct motif families ranging from 14-20 bp. For each motif family, we generated consensus sequences and created position weight matrices (PWMs) for downstream analysis.
3. **Transcription Factor Binding Site Mapping:** We systematically screened all 42 motif PWMs against the JASPAR 2022 vertebrate transcription factor database using TOMTOM, applying stringent statistical thresholds (p-value < 0.001, q-value < 0.05) to identify high-confidence matches to host regulatory elements.
4. **Statistical Validation of Enrichment**:** For each motif-TF pair, we computed local enrichment (within-sequence positional bias) and global enrichment (frequency across pathogenic vs non-pathogenic strains) using permutation testing. Motifs showing both local and global enrichment (p < 0.01) were prioritized for biological interpretation.

This systematic approach identified 42 conserved motifs mapping to 10 distinct vertebrate transcription factors with statistically significant enrichment in pathogenic viral strains (Supplementary Table S3), revealing coordinated viral strategies for host regulatory machinery hijacking rather than isolated mimicry events.

## 5 Data and Code Availability

The HVUE benchmark datasets, training scripts, and complete implementation are publicly available at https://github.com/duttaprat/HViLM. Pre-trained HViLM-base model weights and fine-tuned task-specific variants (HViLM-Patho, HViLM-Tropism, HViLM-R0) are available on Hugging Face at https://huggingface.co/duttaprat/HViLM-base. The VIRION viral genome database used for continued pretraining is accessible at https://virion.verena.org. Interpretability analysis results, including the 42 discovered motifs with transcription factor mappings, are provided at Supplementary Data. All resources are released under the MIT License.

## 6 Conclusion

We present HViLM, the first foundation model specifically designed for comprehensive viral risk assessment through multi-task prediction of pathogenicity, host tropism, and transmissibility. Through viral-specialized continued pre-training on 5 million genomic sequences and parameter-efficient fine-tuning with LoRA, HViLM achieves state-of-the-art performance across the HVUE benchmark (95.32% pathogenicity, 96.25% host tropism, 97.36% transmissibility), substantially outperforming general genomic foundation models. Beyond predictive accuracy, interpretability analysis reveals that HViLM captures genuine biological mechanisms underlying coronavirus pathogenicity through attention-guided discovery of host transcription factor binding site mimicry. The identification of 42 conserved motifs matching 10 distinct vertebrate transcription factors—with convergent evolution of 8 independent sequences targeting Irf1 for immune evasion and multiple motifs targeting Foxq1 for epithelial tropism—demonstrates that pathogenic viruses employ coordinated multi-target genomic strategies to hijack host regulatory machinery. These findings establish foundation models as powerful tools not only for rapid computational characterization of emerging viral threats but also for mechanistic discovery of pathogenicity determinants, supporting both pandemic preparedness through scalable risk assessment and therapeutic development through identification of candidate antiviral targets. The released HVUE benchmark, pre-trained model weights, and evaluation protocols provide the research community with standardized resources to advance viral genomics foundation models and accelerate preparedness for future pandemic threats.

